# Thermodynamics of Homopeptide Aggregation

**DOI:** 10.1101/2020.01.27.921700

**Authors:** Tam T. M. Phan, Jeremy D. Schmit

## Abstract

Amyloid aggregates are found in many neurodegenerative diseases including Huntington’s, Alzheimer’s, and prion diseases. The precise role of the aggregates in disease progression has been difficult to elucidate due to the diversity of aggregated states they can adopt. Here we study the formation of fibrils and oligomers by exon 1 of huntingtin protein. We show that the oligomer states are consistent with polymer micelles that are limited in size by the stretching entropy of the polyglutamine region. The model shows how the sequences flanking the amyloid core modulate aggregation behavior. The N17 region promotes aggregation through weakly attractive interactions, while the C38 tail opposes aggregation via steric repulsion. We also show that the energetics of cross-*β* stacking by polyglutamine would produce fibrils with many alignment defects, but minor perturbations from the flanking sequences are sufficient to reduce the defects to the level observed in experiment. We conclude with a discussion of the implications of this model for other amyloid forming molecules.

## INTRODUCTION

Protein aggregates are implicated as the causative factor in numerous diseases, including neurodegenerative diseases like Alzheimer’s and Huntington’s (1, 2). The most conspicuous of these assemblies are insoluble fibrils consisting of molecules stacked in a cross-*β* motif. However, the predominant evidence is that disease progression is actually driven by smaller, soluble oligomers (3). These states are more difficult to study than fibrils because they tend to be transient and heterogeneous. In most cases it is believed that the oligomers are metastable with respect to the fibril, but favored kinetically due to the fact that they lack the large nucleation barrier associated with fibril formation (4–10). Confounding the issue is the fact that in *vitro* conditions inevitably differ from those *in vivo*, raising the question of whether the oligomers observed in the lab are the same as those occurring naturally. This question would be more readily answered with an understanding of the nature and stability of the various states.

The common features of amyloid diseases give rise to another question. To what extent is aggregation and toxicity dependent on the specific sequence and structural states of the proteins? An interesting case study for this question is exon 1 of huntingtin protein, which contains a polyglutamine core flanked by short, unstructured sequences at the N- and C-terminal ends. The aggregation behavior is driven by the polyQ core, with increasing polyQ lengths correlating with earlier disease onset (11, 12). However, the terminal sequences modulate the aggregation propensity with the N-terminus promoting aggregation and the C-terminus promoting higher solubility (13). The behavior of the latter sequence is not surprising as the C-terminal fragment is composed primarily of proline residues. However, the aggregation promoting property of the N-terminus is more difficult to understand as this segment has a high solubility in isolation [Rohit Pappu, personal communication].

Huntingtin (Htt) shows qualitatively similar aggregation behavior to other amyloid proteins with distinct fibril and oligomer states. The low sequence complexity of huntingtin suggests that these states are not due to sequence-specific interactions, but arise more generally from the polymer nature of the molecule. Here we show that the stability of these states can be modeled by treating huntingtin as a triblock copolymer.

For a simple polymer, we expect two limiting behaviors; either the swollen random walk of a polymer in good solvent, or the collapsed state typical of a polymer in poor solvent. Recent experiments and simulations have shown that monomeric huntingtin adopts conformations consistent with the poor solvent case (14). Accordingly, we allow the collapsed globules in our model to coalesce further to form copolymer micelles, which we associate with the oligomer state. To account for the fibril state we add a second form of intermolecular interactions in which the backbones and sidechains pack more efficiently at the cost of conformational entropy. Surprisingly, experiments have shown that huntingtin fibrils are highly ordered despite the discrete translational symmetry of the polyglutamine core (15). We show that this alignment specificity arises naturally from the energetics of the binding ensemble, and it is further assisted by the N- and C-terminal regions.

## MODEL

### Monomer and oligomer are modeled as collapsed globule

To construct our free energy for Htt, we take the reference state to be a well-solvated Flory coil. In this state, contacts between amino acids are negligible and the random walk entropy is maximized. This state is purely hypothetical because experiments and simulation have shown that Htt adopts configurations consistent with a polymer in poor solvent (16–19). This means that favorable interactions between amino acids are sufficient to pay the entropy cost to collapse the random coil into a globule. These same interactions can also drive the condensation of Htt molecules into oligomers. Since monomer collapse and oligomer formation are driven by the same desolvation reaction, we describe them both by the free energy

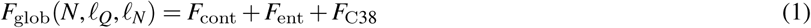

where the terms represent the amino acid contact energy, the change in conformational entropy, and the contribution from the C38 tail. *N* is the size of oligomer in monomer units and *ℓ*_*Q/N*_ are the number of amino acids in the polyQ and N-terminal (if present) regions. The contact energy has a bulk and surface term

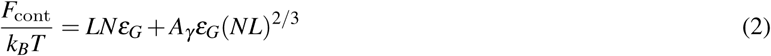

Simulations have shown that the collapsed globule contains both the polyQ region and the N-terminal region (14, 16). Therefore the bulk term is proportional to the total length of these regions *L* = *ℓ*_*Q*_ + *ℓ*_*N*_ where *ℓ*_*N*_ = 17 for molecules containing the N-terminal segment and *ℓ*_*N*_ = 0 for molecules without the tail. For the burial energy we take a weighted average for the desolvation of glutamine and N17 amino acids.

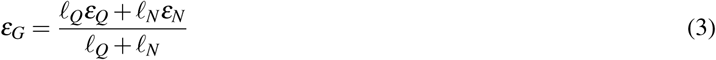

The bulk term, *LNε*_*G*_, over counts the driving force for collapse because residues at the surface of the globule are incompletely desolvated. This is corrected by the surface term *A*_*γ*_ *ε*_*G*_(*NL*)^2*/*3^. Simple geometrical considerations give a value *A*_*γ*_ = − 2.4 for the constant (see Appendix).

The entropic contribution to the free energy takes the form

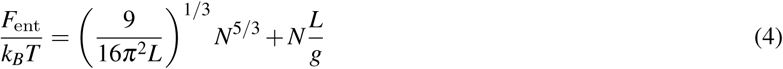

The two terms account for polymer stretching in the oligomer state and the compression of the coils in the collapsed monomer state, respectively. In practice, only one of these terms is significant at any time, which allows for the additive approximation in Eq. 4.

The stretching term provides a repulsive energy that arrests oligomer growth. This term arises in polymer micelles because the molecules must extend from the surface to fill the interior of the aggregate when its radius grows longer than the radius of gyration of a random walk polymer (20). To compute this term we note that the stretching energy for one monomer in the oligomer is 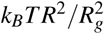 where *R* is the oligomer radius (calculated in the Appendix) and 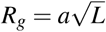 is the radius of gyration for a monomer. Here we have taken the Flory exponent to be *ν* = 1*/*2 since the excluded volume swelling of the polymer will be screened by excluded volume interactions with neighboring molecules. The total stretching energy for the whole oligomer is then

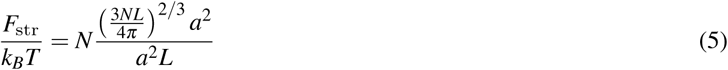

which simplifies to the first term in Eq. 4.

In the monomer state, the molecules face the opposite problem where the entropic loss is due to compression of the random coil. For a polymer under confinement, the free energy change can be estimated by the blob model (21). In this model, the polymer can be sub-divided into statistically independent segments that are each small enough that the effects of confinement are not felt. Confinement effects arise at the interface between these statistical blobs, where it exerts a perturbation on the order of *k*_*B*_*T*. Therefore the free energy of confinement is ∼ *L/*(*gℓ*_*k*_) where *g* is the number of statistically independent segments per blob, and *ℓ*_*k*_ is the Kuhn length. The number of segments per blob can be found by requiring that the segment density per blob *g/*(*ag*^*ν*^)^3^ is equal to the density of the entire system *L/V* where *V* is confinement volume. Therefore *g*^3*ν*−1^ = *V/a*^3^*L*. In this case, we are interested in a collapsed globule where the confinement volume is equal to the total volume of the chain *V* = *La*^3^. This gives *g* = 1. Therefore the compression free energy is just *k*_*B*_*T* times the number of Kuhn lengths. To estimate this, we note that the persistence length of polyQ is about 13 (22) and that the Kuhn length is twice the persistence length (23). Therefore, the statistical correlation along a polymer extends over *ℓ*_*k*_ ≃ 26/3 ≃ 8.7 amino acids.

The final contribution to the globule free energy comes from the C-terminal tail. This region has the sequence P_11_-QLPQPPPQAQPLLPQPQ-P_10_. Given the limited flexibility of proline and the propensity to form polyproline helices, this tail will be more rigid, although largely disordered (24). We assume that the tails interact primarily by excluded volume interactions. Due to the non-uniform flexibility of the tails, it is difficult to apply the blob model to compute the confinement effect due to neighboring tails. Still, inspection of the sequence suggests that 1-3 blobs per tail is reasonable. In fact, our results are insensitive to values in this range. Here we report results for *f*_C38_ = 2 *k*_*B*_*T*.

### The fibril state is a cross-*β* core with disordered tails

Htt fibrils consist of a cross-*β* core that spans the polyQ region but does not include N17 or C38 (15, 24). Evidence suggests that the *β*-sheet core is most likely anti-parallel as shown in Fig. 1 (25), although parallel cores may also occur (26). The specifics of parallel or anti-parallel do not enter our model since the parameters give the average interaction experienced by each sequence block.

**Figure 1:**
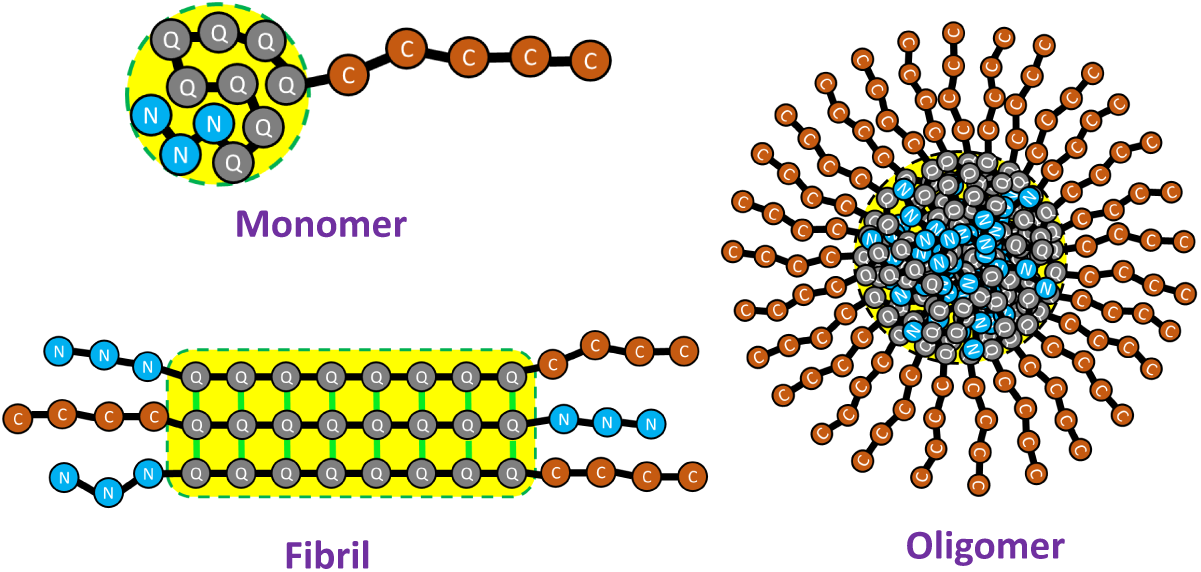
Cartoon representation of the three states of Htt. In the monomer state, the peptide collapses into a globule containing both polyQ and N-terminal regions. The oligomer state is a micelle-like assembly of a few thousand monomers with a spherical core containing the polyQ and N- terminal regions. The fibril state is a cross-*β* amyloid core of polyglutamine flanked by disordered tails on both sides.

We write the fibril free energy as a *β*-sheet term that scales linearly with the length of the polyQ core, modified by perturbations from the terminal segments.

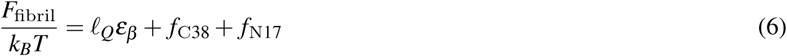

Here *ε*_*β*_ is the free energy gain to move one glutamine residue from the solvated random coil state into the cross-*β* core. The second term accounts for the interaction between C-terminal segments which we expect to be the same as in the globule state. Finally, *f*_N17_ accounts for the interaction of N17 tails. These segments are soluble, but not purely repulsive like the proline-rich C38 (15). To account for the possibility of sequence specific interactions between N17 tails, we obtain this parameter by fitting.

### Critical concentrations are computed from the change in free energy

The free energies derived in the previous sections can be compared to the critical concentration measured in experiments. We start by writing down the equilibrium constant for N-fold oligomerization.

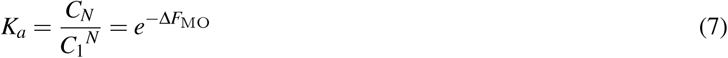

where

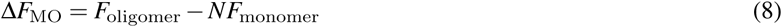

We identify the critical concentration for oligomerization as the point where there is an equal amount of protein in the monomer and oligomer states *C*_1_ = *NC*_*N*_, which can be combined with Eq. 7 to yield a relationship between the critical concentration and the free energy of the oligomer state

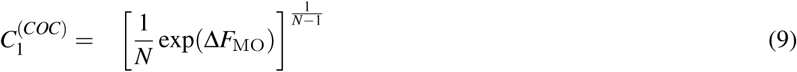

Eq. 9 requires the size of the oligomer, *N*, which we obtain by minimizing Eq. 8

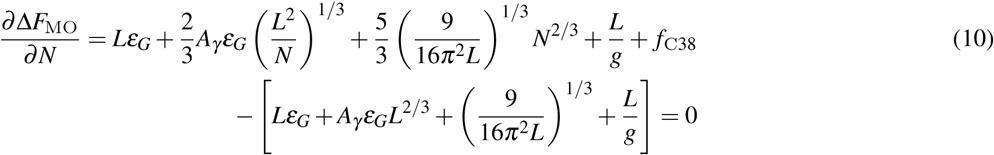

The formation of fibrils can also be associated with a critical concentration. However, unlike the soft transition seen in oligomers, the critical concentration for fibril formation is very sharp (27). This is analogous to the solubility limit in a phase transition. The critical concentration for fibril formation is (27)

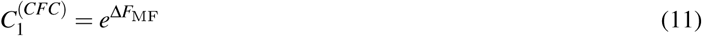

where

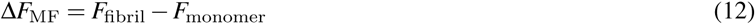

is the free energy for transferring a molecule from the monomer state to the fibril state.

### Fibril alignment defects incur a free energy penalty

Atomic resolution models of amyloid fibrils show striking order in the alignment of molecules (28). However, it is not known whether this order in generally present or if it is an artifact of structural methods that are limited to systems that possess such order. PolyQ aggregates represent an extreme test of the alignment tendency of amyloids due to the discrete translational symmetry they possess.

Here we introduce an equilibrium model to compute the frequency of alignment defects in polyQ fibrils. Following previous work (29), we quantify the alignment using the registry variable *R*, which can take the values − *ℓ*_*Q*_ < *R* < +*ℓ*_*Q*_, where *ℓ*_*Q*_ is the number of glutamine residues in each molecule. The value *R* = 0 denotes the in-register state, positive values of *R* indicate that the incoming molecule is shifted toward its C-term, while negative values of *R* indicate a shift to the N-term (see Fig. 2).

**Figure 2:**
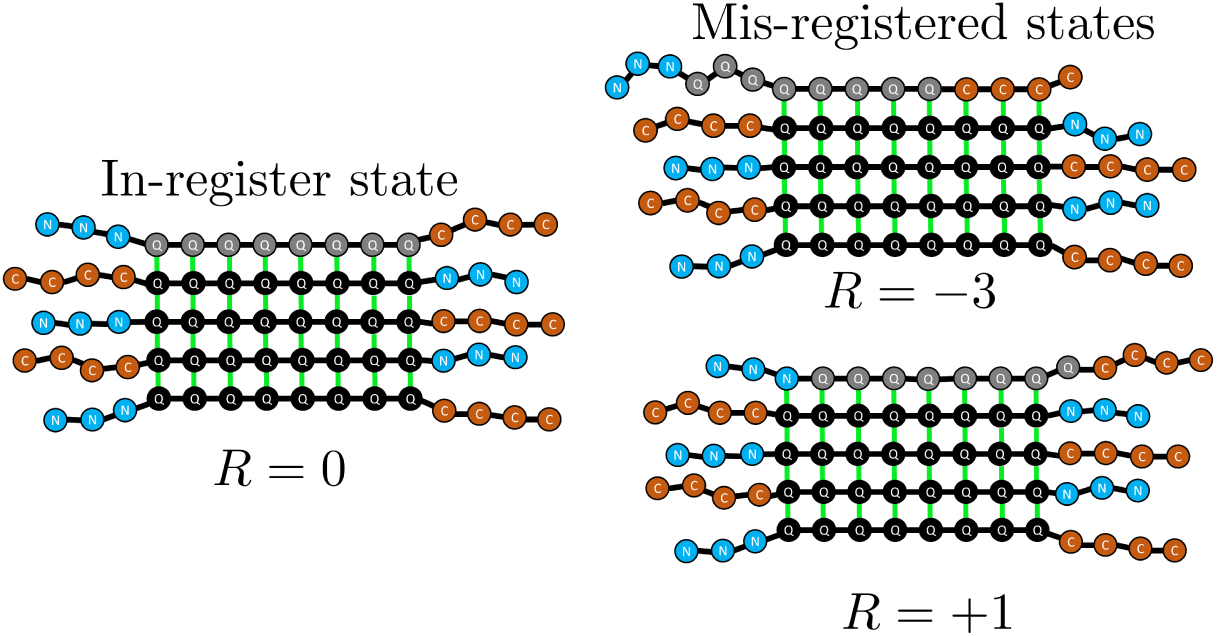
Cartoon representation of the in-register state and mis-registered states. The registry variable, *R*, defines the alignment of an incoming molecule with the existing fibril. *R* = 0 indicates perfect alignment of the polyQ region, while negative and positive values indicate N-terminal and C-terminal shifts, respectively.

For mis-aligned states with *R* < 0, there will be H-bonds between glutamines of the existing fibril and amino acids in C38 of the incoming molecule. Conversely, if *R* > 0, there will be H-bonds between glutamines of the existing fibril and N17 of the incoming molecule.

We compute the probability for a given alignment by

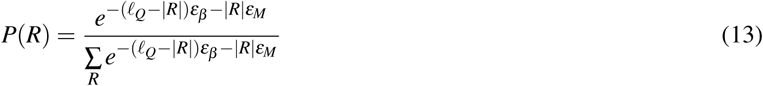

where the denominator is the partition function for the alignment states and *ε*_*M*_ is equal to *ε*_NQ_ or *ε*_CQ_ to account for interaction between the polyQ core and the N- or C-terminal tail of mis-aligned molecules. The mis-alignment energy is not symmetric because we assume that residues lying outside the *β*-core are too disordered to have a significant interaction energy.

## RESULTS

### Polyglutamine desolvation competes with polymer entropy

To obtain values for the energies appearing in the model we fit the calculated critical concentration to the experiments of Crick et al (13). There are four free parameters: *ε*_*Q*_, *ε*_*N*_, *ε*_*β*_, and *f*_N17_. Before fitting it is necessary to convert the experimental concentrations to the dimensionless concentration units required by the theory. We do this using a lattice approximation in which the lattice size is set by the size of a water molecule *C*_th_ = *C/*55.5 M. Other choices for rescaling the concentration would result in a constant shift to the free energy which would not affect the results in a meaningful way. The measured and fitted free energy are compared in Fig. 3. The agreement is good with discrepancies ranging from 0.2-1.1 *k*_*B*_*T*.

**Figure 3:**
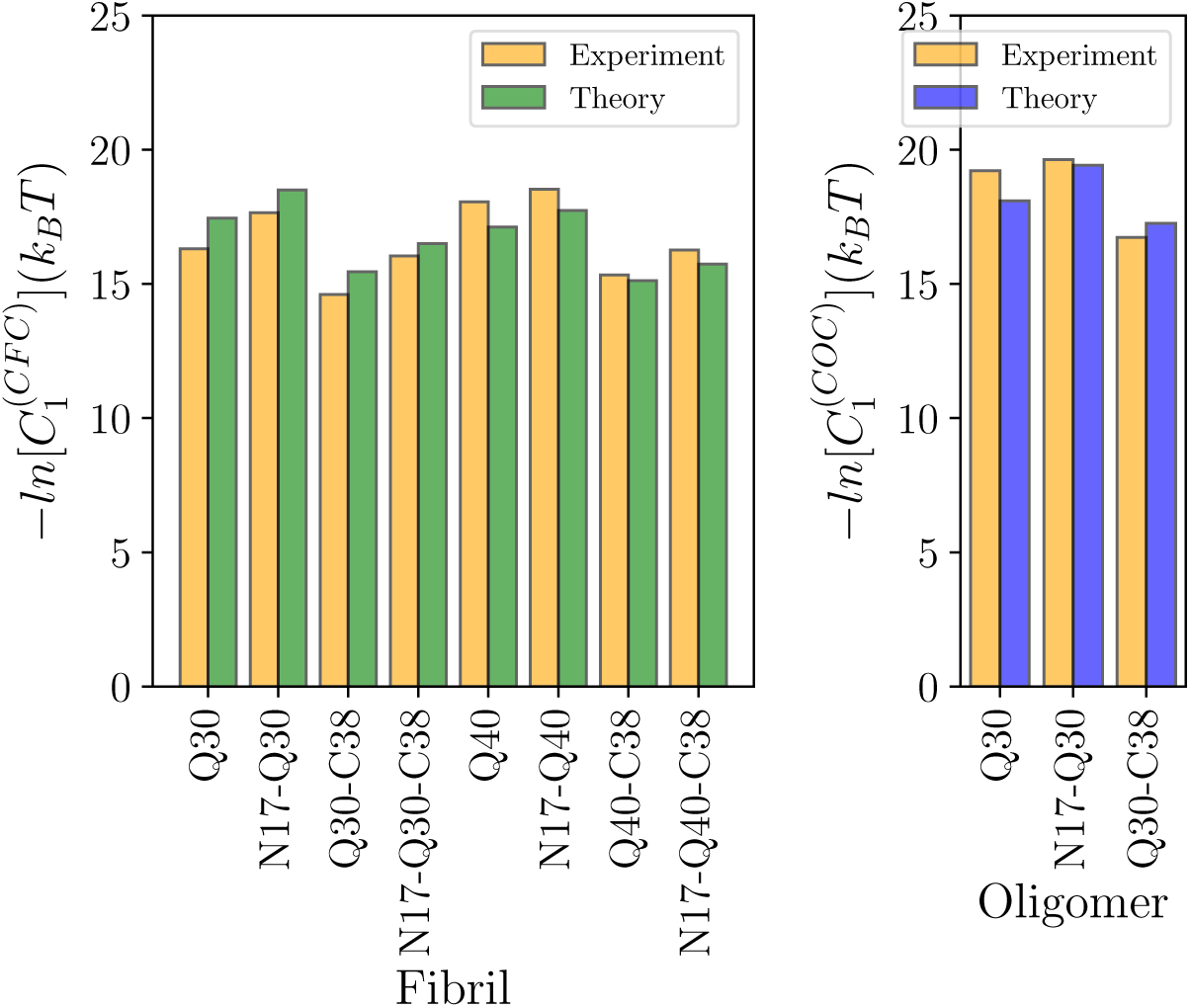
Comparison between the theoretical model and experimentally measured critical concentrations. The model captures the effects of N17 and increasing polyQ length in promoting aggregation and the effect of C38 in inhibiting it.

The parameter values, shown in Table 1, help to clarify the driving forces for aggregation. The free energy of *β*-sheet formation is almost 1 *k*_*B*_*T*, which is stronger than the ∼ 0.5 *k*_*B*_*T* found for A*β* and other amyloid forming molecules (27, 30). This is likely due to fact that polyQ is a homopolymer where all amino acids contribute equally, while other molecules have sequence heterogeneity as well as portions of the molecule in hairpins and disordered fragments that do not contribute to the stability. Interestingly, we find that the free energy of glutamine burial in the oligomer state, *ε*_*Q*_, is even stronger than that of *β*-sheet formation. This reflect the fact that Htt is one of the few molecules where the oligomer state has a lower critical concentration than the fibril state (13, 17). However, it should also be noted that the entropic penalty for elongating the peptide into a *β*-strand is included in *ε*_*β*_, while the conformational entropy contributions to globule formation are separately calculated in Eq. 4.

**Table 1:** Parameters obtained by model fitting

| Parameter | Value |
| --- | --- |
| $\epsilon_Q$ | $-2.14 k_B T$ |
| $\epsilon_N$ | $-0.88 k_B T$ |
| $\epsilon_\beta$ | $-0.94 k_B T$ |
| $f_{N17}$ | $-11.00 k_B T$ |

The model allows us to understand several features of the aggregation behavior. From our results, the oligomer sizes are in the range of 3300-4900 monomers with radii of 9-11 nm for peptides with polyglutamine length of 30. These are in good agreement with the reported experimental size of roughly 10 nm (13).

Fig. 4 shows the free energy of monomer collapse as a function of the polyQ length. In the absence of the N17 tail our model predicts that the free energy is zero for *ℓ*_*Q*_ = 17, meaning that peptides with fewer glutamines will be found primarily in the expanded state while longer polyQ regions will favor the collapsed state. In the presence of N17 the crossover point is at *ℓ*_*Q*_ = 3. While this is fewer glutamine residues than molecules without N17, the total peptide length is longer (20 vs. 17 amino acids) reflecting the fact that it takes more N17 residues to achieve the same desolvation energy of the glutamines.

**Figure 4:**
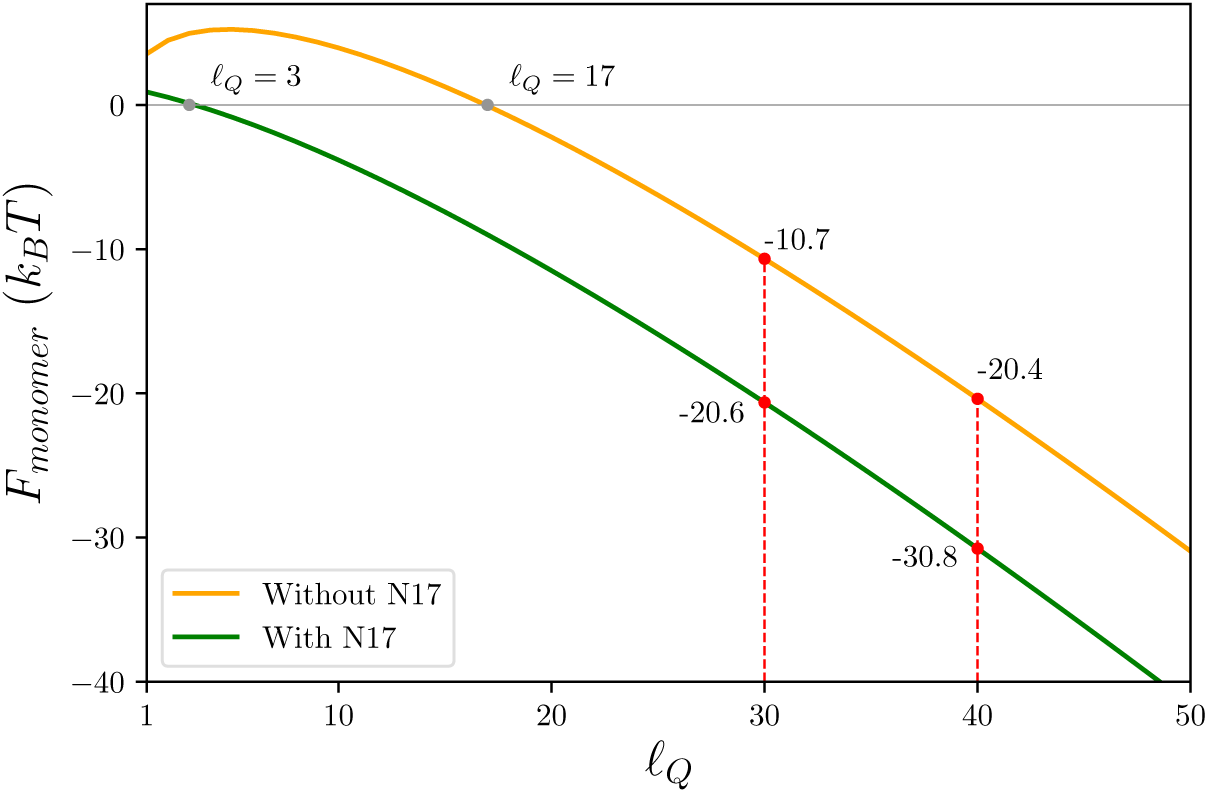
Predicted free energy of monomer collapse for peptides with and without the N-terminal tail as a function of *ℓ*_*Q*_. The results show that peptides with fewer glutamines will prefer the expanded state while longer glutamine peptides will favor the collapsed states. The presence of the N17 tail contributes to the collapse free energy, but less strongly than glutamine residues.

Fig. 5 shows the oligomer free energy as a function of polyQ length in the presence and absence of the flanking regions. Increasing the peptide length, either by adding glutamines or N17 residues, results in larger oligomers because the longer molecules can more easily stretch to fill the interior of the oligomer. However the C38 region adds a repulsive energy that favors smaller oligomers.

**Figure 5:**
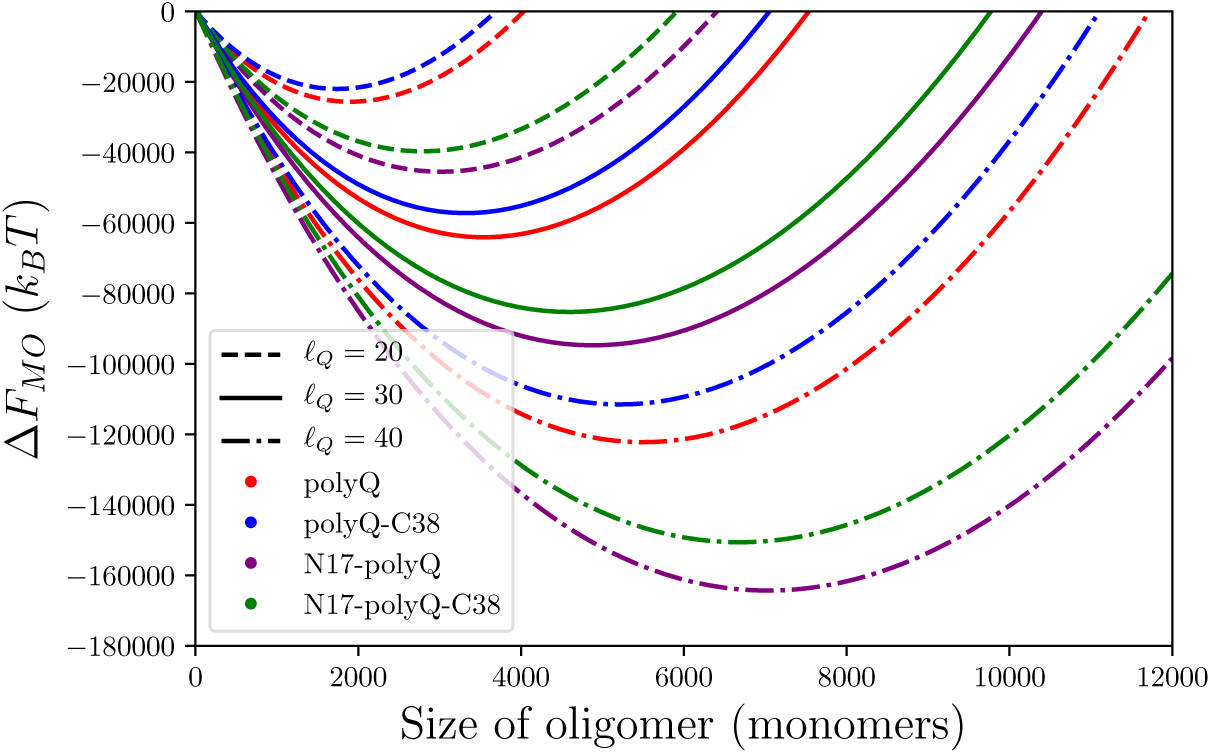
Predicted free energy of oligomer formation for *ℓ*_*Q*_ = 20, 30 and 40 in the presence and absence of N- and C-terminal tails. Increasing the length of the polyglutamine region or adding the N17 tail results in larger oligomers because the extra length more easily stretches to fill the oligomer core. However, adding the C38 tail adds a repulsive energy that favors smaller oligomers.

### Flanking sequences prevent large alignment errors

The parameter *ε*_*Q*_, obtained by fitting the fibril solubilities, can also be used to compute the frequency of registry errors in polyQ fibrils. NMR experiments have shown that single amino acids shifts occur at frequency of 25% (*R* = +1) and 15% (*R* = −1), with larger shifts occurring below the detection limit (15). In comparison, a simple version of our model that does not account for the N- and C-terminal tails (Eq. 13 with *ε*_NQ_ = *ε*_CQ_ = 0) yields registry errors of 17% for *R* = ±1 and 7% for *R* = ±2 (blue line, Fig. 6). From this we make two observations. First, even the weak 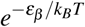 penalty for registry shifts is sufficient to prevent registry errors for most molecules. Secondly, the presence of the N- and C-terminal tails have the dual effect of suppressing shifts of |*R*| > 1 and breaking the symmetry between the shift directions. An inspection of the sequence readily suggests mechanisms by which this may occur. The C-terminus of the polyQ region is connected to a stretch of 28 prolines. Prolines will be poorly tolerated in the cross-*β* core due to their lack of a backbone H-bond donor and their inability to adopt the extended *β*-sheet conformation. To account for this we add a free energy penalty for negative registry shifts. Fig. 6 shows that *ε*_CQ_ values between 0.25 and 1.0 *k*_*B*_*T* have the expected effect of shifting the alignment distribution closer to the experimental observation. However, they also raise the probability of *R* = +2 shifts above 10%, which would be observed experimentally. This discrepancy is easily resolved by an inspection of the N-terminal tail, which has a sequence MATLEKLMKAFESLKSF, with the serine and phenylalanine incorporated in the cross-*β* core (15). This means that positive registry shifts would move the lysine into the cross-*β* core. While the long sidechain could presumably allow for partial solvation of the charge for a *R* = +1 shift, larger registry shifts would require a total desolvation of the charge, thereby incurring a large free energy penalty. If we exclude registry shifts larger than *R* = +1, the predicted distribution of registries is in nearly perfect agreement with experiment (Fig. 6, inset).

**Figure 6:**
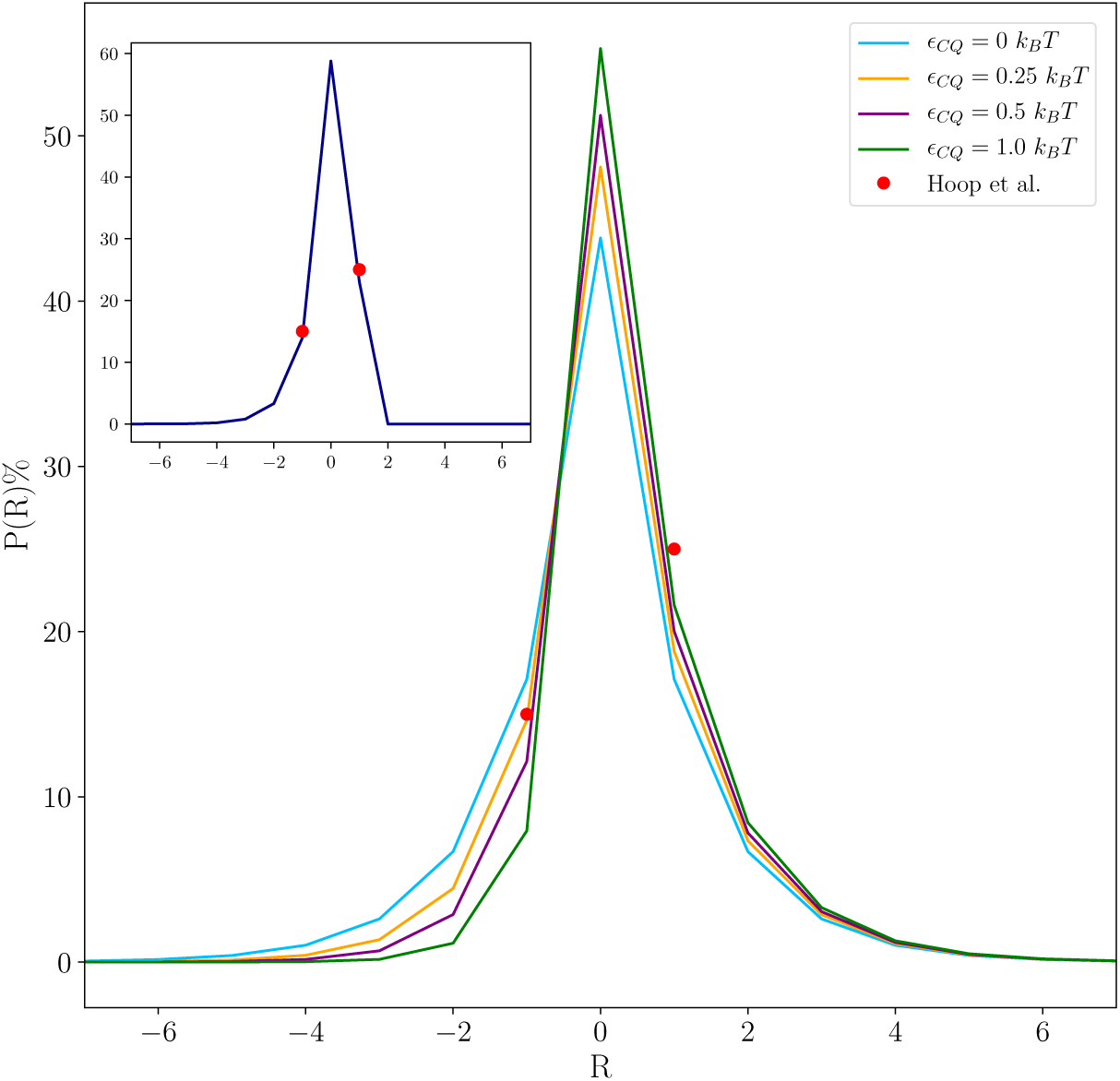
Probabilities of mis-aligned molecules within an Htt fibril as a function of the alignment registry *R* and *ε*_CQ_ (for *ε*_*β*_ = −0.94 *k*_*B*_*T*). The inset shows alignment probabilities for *ε*_CQ_ = 0.5 *k*_*B*_*T* with an additional constraint preventing states with *R* > 1, since this would lead to the burial of the lysine charge.

## DISCUSSION

The micelle-like oligomers described by our theory contrast with the highly ordered *β*-barrel oligomers that have been reported for other amyloid forming molecules (31, 32). It is difficult to imagine a low complexity sequence like huntingtin adopting such an ordered state. But it is worth asking whether the micellar structure of huntingtin might also be formed by other molecules. Supporting this view is the fact that the A11 antibody, which specifically recognizes amyloid oligomers, was developed by forcing A*β* into a micelle-like structure (33). In addition, hydrophobicity correlates strongly with aggregation propensity (34, 35), suggesting that most amyloid-forming moleules will contain a stretch of hydrophobic amino acids sufficiently long to form a polymer micelle.

Our model also provides insights into the mechanism of fibril formation. Specifically, there is the question of whether the highly ordered fibrils reported from NMR or X-ray studies are typical or an artifact of structural methods that work best with ordered systems. Our results show that even for a homopolymer, the binding energy is sufficient to align almost half of the molecules. Also, consistent with previous work, only minor perturbations from a uniform sequence are necessary to generate highly ordered fibrils (29). In the equilibrium analysis employed in this work, the fraction of alignment defects is independent of peptide length. However, the kinetic search over alignments scales exponentially with the peptide length, meaning longer peptides will be more easily trapped in non-equilibrium states under conditions of rapid aggregation (27, 29).

In conclusion, we have shown that block copolymer model is able to explain many features of oligomer and fibril formation in huntingtin. These findings may also have broader implications for other amyloid forming systems.

## Appendix

### Calculation of surface constant

The bulk energy term of Eq. 1 accounts for the desolvation of every amino acid in the globule. However, amino acids on the surface of the globule will only be partially desolvated. To estimate the surface correction to the desolvation energy, we assume that amino acids on the surface only get half the desolvation energy. This gives

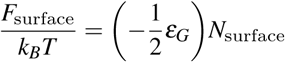

where *N*_surface_ is the number of amino acids on the surface of the globule.

To calculate the *N*_surface_, we relate the radius of the globule to the number of molecules

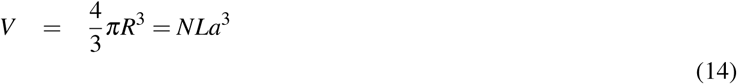

where *a*^3^ is the volume of an amino acid. The number of residues on the surface is

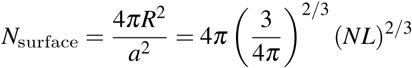

The surface term is therefore

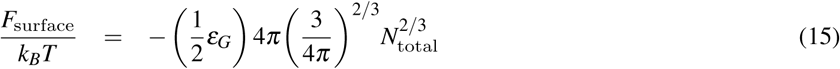

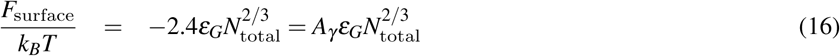

## ACKNOWLEDGMENTS

This work was supported by National Institutes of Health grant R01GM107487.

